# Engineered CTLA-4 targeting chimeras for targeted protein degradation and immune synapse reprogramming

**DOI:** 10.64898/2026.09.29.755403

**Authors:** Ping Ren, Shao-Yu Fang, Shushu Zhao, Hanbing Cao, Kaiyuan Tang, Xingbo Shang, Cole W. Christopher, Yunfei Jiao, Medha Majety, Andre Levchenko, David B. Weiner, Sidi Chen, Xiaoyu Zhou

## Abstract

Despite the success of immune checkpoint blockade in cancer therapy, overcoming the immunosuppressive tumor microenvironment remains a major barrier to durable responses. Here, we introduce cytotoxic T-lymphocyte-associated antigen-4 (CTLA-4)-targeting chimeras (cTACs), a class of synthetic molecules that harness CTLA-4-mediated trans-endocytosis (TE) for targeted protein degradation across cellular boundaries. The lead candidate, cTAC4.0, is composed of the extracellular domain of CD80 fused to an anti-programmed death-ligand 1 (PD-L1) fragment via a human immunoglobulin G4 (IgG4) heavy chain. Unlike existing protein degraders that function within cancer cells via house-keeping pathways, cTAC4.0 operates *in trans* between immune and tumor cells. It redirects the natural CD80-CTLA-4 internalization to capture and degrade PD-L1 from tumor cells into T cells, while simultaneously converting PD-L1-mediated suppression into CD28-driven co-stimulation, depending on the relative availability of CTLA-4 or CD28. Mechanistically, we demonstrate that cTAC4.0 promotes CTLA-4-dependent internalization and lysosomal degradation of both membrane-bound and soluble PD-L1, reduces surface PD-L1 levels on target cells, and activates T cells in a CD28-dependent manner. Functionally, cTAC4.0 enhances the cytotoxicity of both bispecific T cell engagers and T cell receptor (TCR)-engineered T cells *in vitro,* and when combined with these engineered T cells, it suppresses tumor growth *in vivo* in a glioblastoma xenograft model. By coupling immune checkpoint degradation with T cell activation in an immune synapse-enriched manner, cTACs establish a paradigm for intercellular protein degradation systems that simultaneously deplete suppressive ligands and enhance T cell function to overcome resistance in cancer immunotherapy.

## Introduction

Immune checkpoint blockade targeting PD-1 ^1^, PD-L1 ^2^, or CTLA-4 ^3, 4^, has revolutionized cancer immunotherapy by achieving long-lasting responses in a subset of patients. However, most patients either fail to respond initially or eventually develop acquired resistance ^5, 6^. A recurring obstacle is the persistence of immunosuppressive ligands at the tumor-immune interface. Even in the presence of antibody-mediated receptor blockade, these ligands remain abundant on tumor and stromal cells, sustaining a suppressive microenvironment ^7, 8^. Soluble forms of these ligands further contribute to systemic immunosuppression ^9, 10^. Therefore, strategies that physically eliminating suppressive ligands from the immune synapse, moving beyond mere receptor blockade, are urgently needed.

Small molecule-based degraders, such as proteolysis-targeting chimeras (PROTACs) ^11^, offer advantages over antibody blockade by enabling elimination rather than mere inhibition of target proteins. Yet designing high-affinity small molecules for many therapeutically relevant targets remains challenging. Recently, large molecular degraders targeting endocytic receptors, such as TransTACs (utilizing transferrin receptor) ^12^ and EndoTag (utilizing insulin-like growth factor 2 receptor) ^13^, have shown great promise in degrading both transmembrane and soluble proteins in cancer cells. Nevertheless, existing degraders generally lack contextual specificity. They act broadly across cell types, raising concerns about on-target depletion in healthy tissues and perturbation of normal cellular functions. Critically, these systems are not integrated with immune recognition or restricted to sites of antigen-specific T cell activation. This limitation underscores the need for a degrader platform that is preferentially active at immune synapses and dynamically coupled to T cell activation states. Such an approach would enable the selective elimination of suppressive ligands precisely where they impair anti-tumor immunity, while minimizing disruption to essential proteins in non-immune tissues.

CTLA-4 biology provides a unique opportunity for such a strategy. CTLA-4 is a coinhibitory receptor expressed predominantly by activated effector T cells and constitutively by regulatory T cells (Tregs). It regulates immunity through trans-endocytosis (TE), a process whereby CTLA-4 captures CD80/CD86 from the surface of antigen-presenting cells (APCs) and internalizes these ligands for lysosomal degradation within the T cell ^14, 15^. This process depletes costimulatory ligands from the immune synapse and functions as a natural, immune-restricted degradation pathway that dampens CD28-mediated activation. Despite its well-established role in immune suppression, the therapeutic potential of repurposing CTLA-4-mediated TE for redirected protein degradation remains unexplored.

Here, we present CTLA-4-targeting chimeras (cTACs), a class of synthetic molecules that leverage CTLA-4-mediated TE for targeted protein degradation across cellular boundaries. To demonstrate this concept, we engineered cTAC4.0 by fusing the extracellular domain of CD80 to an anti-PD-L1 single-chain variable fragment (scFv) via a human IgG4-Fc domain. We demonstrate that cTAC4.0 suppresses native CD80 TE while promoting PD-L1 degradation *in trans* and converts PD-L1-mediated suppression into CD28-driven co-stimulation. Beyond these core mechanisms, we further characterize the functional versatility of cTACs in multiple contexts, including their ability to degrade soluble PD-L1 and enhance the cytotoxicity of both bispecific T cell engagers (BTEs) and TCR-engineered T cells *in vitro* and *in vivo*. By coupling targeted protein degradation with immune regulation at the level of the immune synapse, cTACs establish a paradigm for overcoming ligand-mediated immune suppression in cancer.

## Results

### Development of cTACs to redirect CD80 trans-endocytosis towards PD-L1 degradation

CTLA-4-mediated TE of CD80/CD86 is a well-characterized mechanism of immune suppression^14, 15^. We hypothesized that this process could be repurposed to mediate the TE of co-inhibitory molecules such as PD-L1, a critical mediator of immune evasion in the tumor microenvironment (TME) ^16^(**Fig.1a**). To test this, we engineered multiple cTACs incorporating either the extracellular domain of human CD80 (CD80-ECD) or an scFv derived from tremelimumab ^17^. To engage PD-L1, we designed cTACs with either a high-affinity PD-1 extracellular domain (ha-PD1) ^18^ or an scFv derived from avelumab ^19^ (**Fig. 1b**). To assess their ability to mediate PD-L1 removal, we employed an *in vitro* coculture assay with PD-L1-BFP-overexpressing NALM6 cells as donors and CTLA-4-overexpressing Jurkat T cells as recipients. (**Fig. 1c**). Compared to control groups, all cTACs, except cTAC3.0, induced efficient, dose-dependent PD-L1-BFP TE (**Fig. 1d**). Notably, cTAC4.0 exhibited potent PD-L1 TE at concentrations as low as 0.1 nM (**Fig. 1d, S1a-b**). A slight reduction in TE was observed at 10 nM, likely due to CTLA-4 internalization and degradation, consistent with previous studies showing that enhanced CD80-CTLA-4 interactions can trigger CTLA-4 downregulation ^20^ (**Fig. 1d**). Based on these findings, cTAC4.0 was selected as the lead candidate for further experiments.

**Figure 1.**
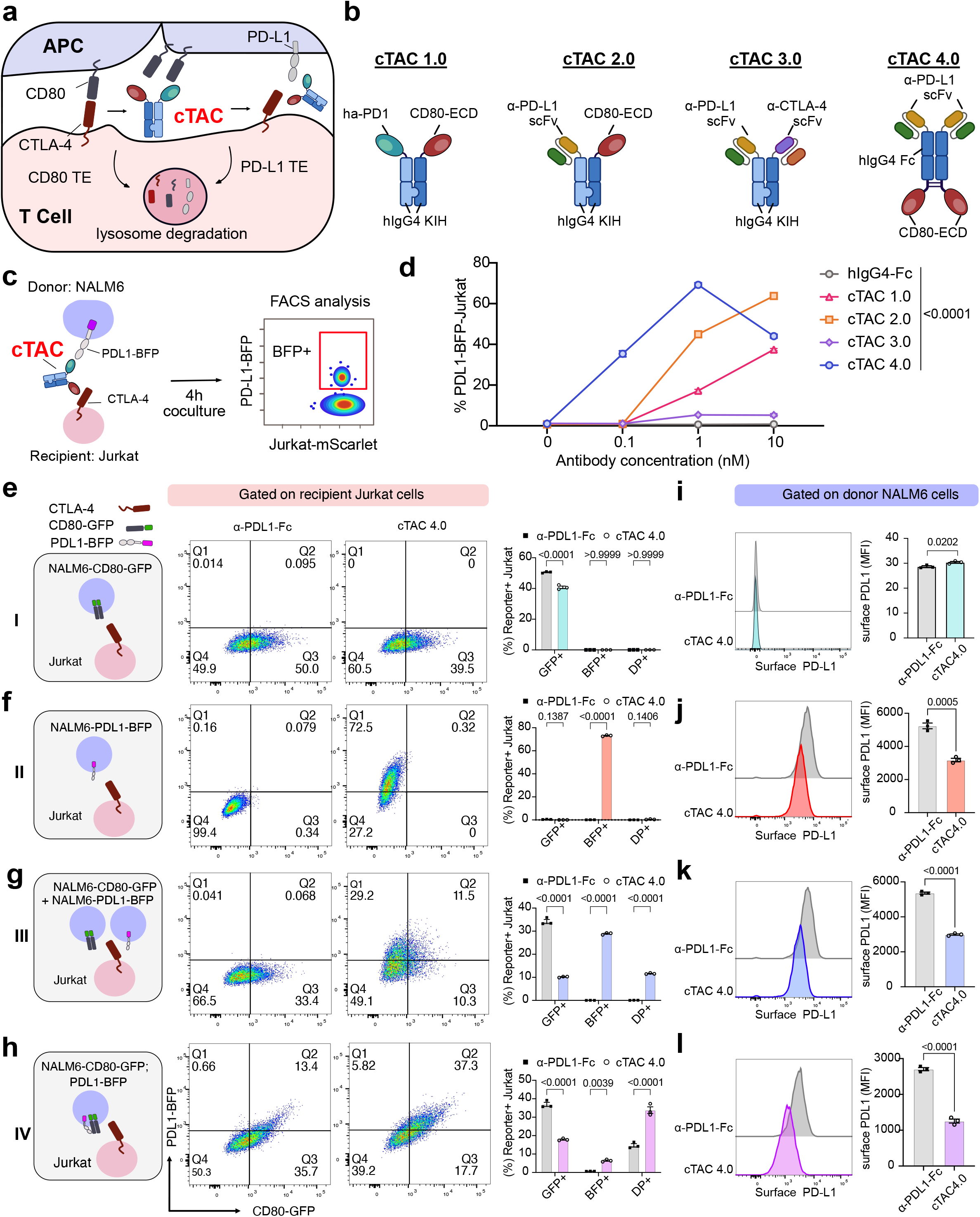
Development of cTACs to redirect trans-endocytosis of CD80 towards PD-L1. **(a)** Schematics demonstrating CD80 trans-endocytosis by CTLA-4, and hypothesized mechanism of action (MOA) of utilizing cTACs to redirect trans-endocytosis of CD80 towards PD-L1 degradation. **(b)** Schematics showing the four versions of cTACs developed for testing. In cTAC1.0, cTAC2.0 and cTAC3.0, a knob-in-hole human IgG4 backbone was used. In cTAC4.0, a human IgG4 heavy chain backbone was used. For the PD-L1 binder, a high affinity PD-1 (ha-PD-1) was used in cTAC1.0; and a scFv derived from avelumab was used in cTAC2.0-4.0. For the CTLA-4 binder, human CD80 extracellular domain (hCD80-ECD) was used in cTAC1.0, cTAC2.0, and cTAC4.0; and an scFv derived from tremelimumab was used in cTAC3.0. **(c)** Schematics of workflow used in **d**. PD-L1 fused with fluorescent protein BFP was expressed on donor NALM6 cells, and CTLA-4-expressing Jurkat cells were used as recipient cells. cTAC mediated PD-L1 TE was quantified by BFP transfer from donor NALM6 cells expressing PD-L1-BFP cells to recipient CTLA-4^+^ Jurkat cells using flow cytometry. **(d)** Quantification of transferred PD-L1-BFP on recipient Jurkat cells mediated by cTACs at various concentrations. **(e-h)** Representative flow cytometry plots (left panel) and quantification (right panel) of transferred CD80-GFP and PD-L1-BFP on recipient Jurkat cells induced by 1nM cTACs. Donor NALM6 cells expressing either CD80-GFP only (**e**), PD-L1-BFP only (**f**), CD80-GFP and PD-L1-BFP on separate cells (**g**), or co-expressed CD80-GFP and PD-L1-BFP (**h**). **(i-l)** Representative flow cytometry plots (left panel) and quantification (right panel) of surface PD-L1 on donor NALM6 cells expressing either CD80-GFP only (**i**), PD-L1-BFP only (**j**), CD80-GFP and PD-L1-BFP on separate cells (**k**), or co-expressed CD80-GFP and PD-L1-BFP (**l**). Data are shown as mean ± s.e.m. One-way ANOVA with Dunnett’s multiple-comparisons test was used to assess significance (right panels in **i-l**). Two-way ANOVA with Tukey’s multiple-comparisons test was used to assess significance (**d**, right panels **e-h**). Exact P values are labeled above all comparisons. All numbers defined by ‘n’ indicate the number of biologically independent samples. Data are representative of two independent experiments.

To confirm that PD-L1 transfer is specifically driven by the integrated configuration of cTAC4.0 rather than simply binding to CTLA-4 and/or PD-L1, we generated additional control antibodies and characterized their biophysical properties (**Fig. S2a-d**). cTAC4.0 exhibited strong binding to both PD-L1 and CTLA-4, whereas anti-PD-L1-Fc and CD80-Fc bound only to PD-L1 and CTLA-4, respectively (**Fig. S2b-d**). IgG4-Fc control showed no binding to either antigen (**Fig. S2b-d**). As expected, both cTAC4.0 and CD80-Fc displayed weak binding to CD28 (**Fig. S2b-d**). In coculture assays analogous to those in **Fig. 1c**, PD-L1 transfer was observed exclusively in the cTAC4.0 group, with no significant uptake in any other control groups (**Fig. S3a-b**). These findings indicate that the integrated configuration of cTAC4.0 to engage PD-L1 and CTLA-4 simultaneously is essential for PD-L1 transfer.

CTLA-4-mediated CD80 TE naturally occurs when T cells acquire CD80 from antigen-presenting cells^14, 15, 20^. To determine whether cTAC4.0-mediated PD-L1 transfer competes with endogenous CD80 TE, we designed four experimental conditions using donor cells expressing: (i) only CD80; (ii) only PD-L1; (iii) both CD80 and PD-L1 on separate cells; (iv) CD80 and PD-L1 co-expressed on the same cell (**Fig. 1e-h**). Across all conditions, cTAC4.0 significantly reduced CD80 TE while enhancing PD-L1 transfer. Notably, in condition (III), cTAC4.0 effectively redirected CD80 TE toward PD-L1 transfer (**Fig. 1g**). In condition (IV), cTAC4.0 markedly increased the percentage of GFP and BFP double positive (DP^+^) recipient Jurkat cells, suggesting co-transfer of PD-L1 and CD80 when co-expressed on the same donor (**Fig. 1h**). This observation is consistent with the reported *cis*-interaction between PD-L1 and CD80 ^21, 22^. To examine whether PD-L1 transfer affected its surface levels on donor cells, we quantified PD-L1 expression on NALM6 cells post-coculture. cTAC4.0 significantly reduced surface PD-L1 levels on donor cells with initially high PD-L1 expression (**Fig. 1i-l**).

A key feature of CD80 TE by CTLA-4 is internalization and subsequent lysosomal degradation of CD80 ^14^. To determine whether cTAC4.0-mediated PD-L1 transfer follows a similar route, we analyzed the subcellular distribution of CD80-GFP and PD-L1-BFP in the recipient cells from the four conditions in **Fig1.e-h**. Both CD80-GFP and PD-L1-BFP exhibited low colocalization with the membrane protein CD45, indicating efficient internalization after transfer (**Fig. 2a-b**). To further test whether PD-L1 was degraded in the lysosome, recipient Jurkat cells were pretreated with bafilomycin A1 (BafA1), an inhibitor of lysosomal degradation^23^. BafA1 treatment led to marked accumulation of transferred PD-L1 in the cTAC4.0 group, but not in the IgG4-Fc control group (**Fig. 2c-d**). These results indicated that the transferred PD-L1 was internalized and subsequently degraded in the lysosome.

**Figure 2.**
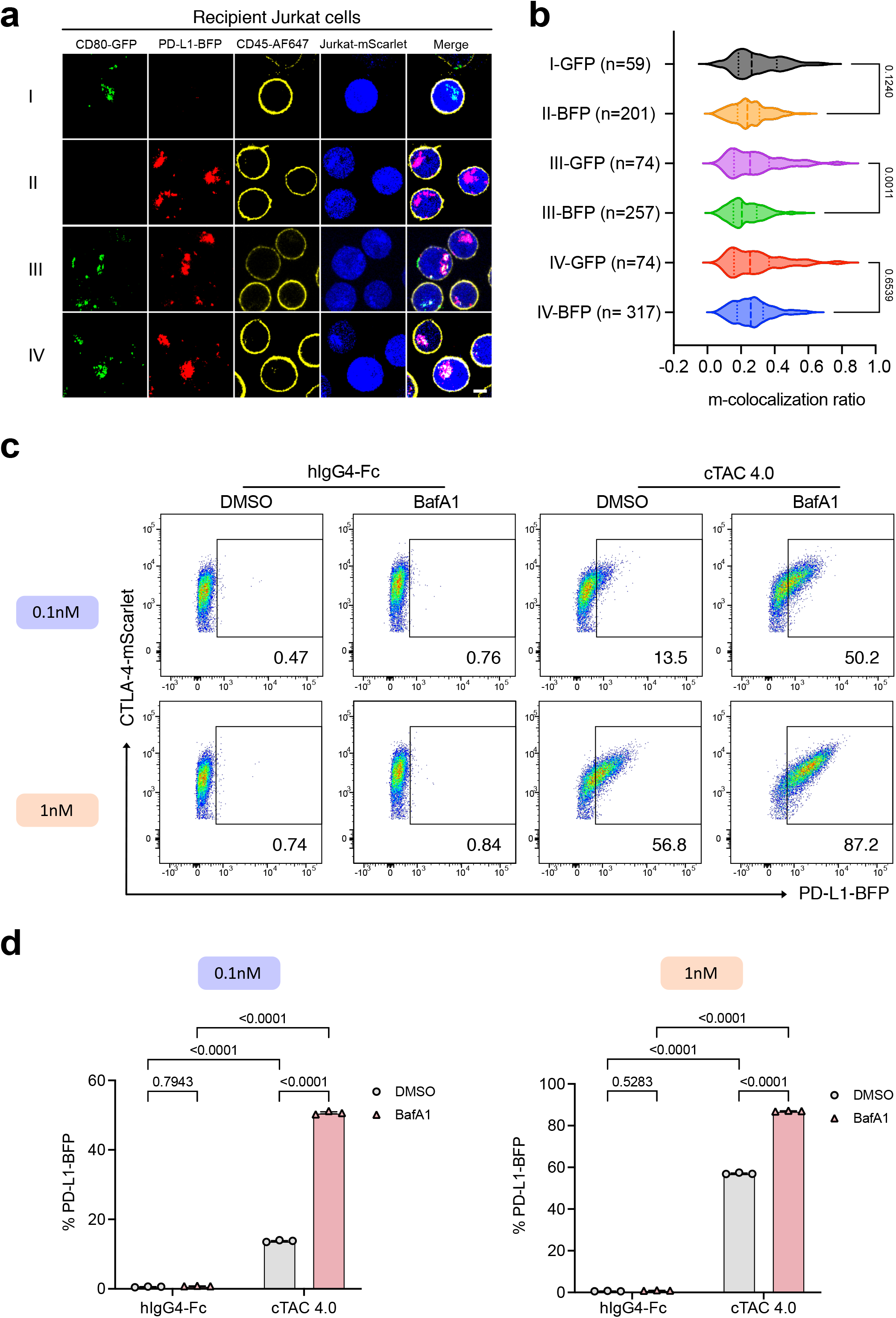
cTAC4.0 mediated the internalization and degradation of transferred PD-L1. **(a)** Representative confocal images, demonstrating the subcellular distribution of transferred CD80-GFP and PD-L1-BFP in recipient Jurkat cells after coculturing with different donor cells as shown in Fig. 1e**-h**. Cell membrane was indicated by surface staining of CD45 using Alexa-fluor 647 labeled antibody (CD45-AF647). **(b)** Quantification of **a**, showing colocalization ratio of GFP or BFP with membrane CD45 signals. In each group, the total number of quantified cells is labeled on the y axis of the plot. **(c)** Representative flow cytometry plots to quantify 0.1nM or 1nM of cTAC4.0 mediated PD-L1 TE by Jurkat-CTLA-4 cells, which were pretreated with either DMSO or BafA1. **(d)** Quantification of **c**. Data are shown as mean ± s.e.m. One-way ANOVA with Dunnett’s multiple-comparisons test was used to assess significance (**b**). Two-way ANOVA with Tukey’s multiple-comparisons test was used to assess significance (**d**). Exact P values are labeled above all comparisons. All numbers defined by ‘n’ indicate the number of biologically independent samples. Data are representative of two independent experiments.

Together, these findings demonstrated the feasibility of repurposing CTLA-4-mediated TE for targeted protein degradation. Specifically, cTAC4.0 facilitated dose-dependent PD-L1 uptake, redirected CD80 TE toward PD-L1 removal, and promoted its lysosomal degradation within recipient cells, while concurrently reducing PD-L1 surface levels on donor cells.

### cTAC4.0 converted PD-L1 mediated suppression to activation

Given that the CD80-ECD incorporated in cTAC4.0 engages not only CTLA-4 but also the costimulatory receptor CD28 (**Fig. S2d**), we next examined whether cTAC4.0 could act as a molecular bridge, simultaneously engaging PD-L1 on donor cells and CD28 on recipient T cells, thereby converting PD-L1-mediated suppression into CD28-driven activation. To test this hypothesis, we co-cultured CTLA-4⁺ Jurkat cells with PD-L1-expressing NALM6 cells. In this setting, only cTAC4.0 significantly activated Jurkat cells in a dose-dependent manner, as indicated by CD69 upregulation, whereas anti-PD-L1-Fc, CD80-Fc, their combination, or IgG4-Fc alone had no effect (**Fig. S4a-b**). This indicates that the activity of cTAC4.0 is not merely due to its ability to bind PD-L1 and CD28 separately, but rather requires its integrated bridging configuration that couples these interactions across cells to promote activation.

Since Jurkat cells expressed minimal endogenous PD-1, we overexpressed either wild-type or truncated PD-1 in Jurkat cell and employed an NF-κB reporter system to assess cTAC4.0-mediated T cell activation under PD-1-mediated suppression (**Fig. 3a**). Regardless of CTLA-4 expression, cTAC4.0 significantly enhanced NF-κB signaling in a dose-dependent manner in Jurkat cells expressing either wild-type PD-1 (PD-1^wt^) or a truncated PD-1 (PD-1^trunc.^) lacking the intracellular domain. In contrast, the controls had no effect (**Fig. 3b-c**). Notably, although activation level was lower in PD-1^wt^ Jurkat cells than in PD-1^trunc.^ cells at low concentrations, cTAC4.0 at 10 nM or higher effectively overcame PD-1-mediated inhibition (**Fig. 3b-c**). These findings indicated that cTAC4.0 can counteract PD-L1-PD-1 suppression and restore Jurkat T cell activation.

**Figure 3.**
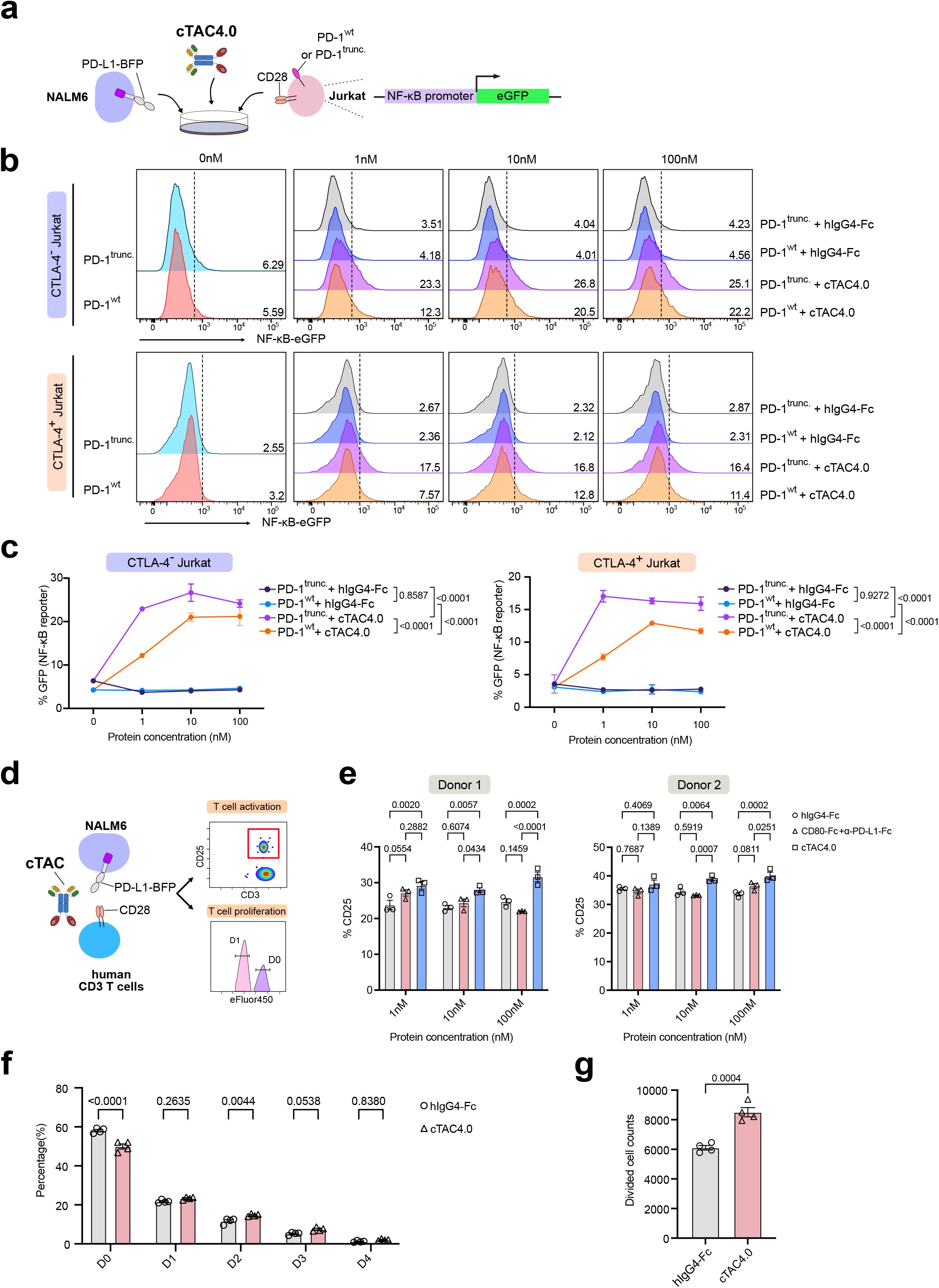
cTAC4.0 converted PD-L1 mediated suppression to T cell activation. **(a)** Experimental schematics for panels **b** and **c**, illustrating the use of cTAC4.0 at various concentration to stimulate CTLA-4^+^ and CTLA-4^−^ Jurkat cells expressing either wild-type PD-1 or a truncated PD-1 lacking the intracellular domain. An NF-κB-eGFP reporter is used to indicate the activation of Jurkat cells in the presence of NALM6-PD-L1-BFP cells. **(b)** Representative flow plots showing the intensity of NFκB-eGFP signals in recipient Jurkat cells with (lower panel) or without (upper panel) expression of CTLA-4. The gray dashed line represents the gating used to determine the percentages of eGFP^+^ Jurkat cells, which are labeled as numbers on the plots. **(c)** Quantification of the percentage of NFκB-eGFP^+^ Jurkat cells under the stimulation of various concentrations of cTAC4.0. **(d)** Experimental schematics for panels **e-g**, demonstrating the use of cTAC4.0 to stimulate primary CD3 T cells in the presence of NALM6-PD-L1-BFP cells pretreated with mitomycin C. T cell activation and proliferation are measured by flow cytometry. **(e)** Quantification of T cell activation induced by varying concentrations of cTAC4.0 in CD3 T cells from two healthy donors, assessed by CD25 expression levels. **(f)** Quantification of the percentage of dividing T cells based on the dilution of the proliferation dye eFluor450. The number labeled on the x-axis indicates the division number modeled with FlowJo. **(g)** Quantification of the number of divided T cells, based on the dilution of the proliferation dye eFluor450, 3 days after *in vitro* stimulation with cTAC4.0. Data are shown as mean ± s.e.m. One-way ANOVA with Dunnett’s multiple-comparisons test was used to assess significance (**f, g**). Two-way ANOVA with Tukey’s multiple-comparisons test was used to assess significance (**c, e**). Exact P values are labeled above all comparisons. All numbers defined by ‘n’ indicate the number of biologically independent samples. Data are representative of three independent experiments.

Next, we explored whether cTAC4.0 could similarly enhance activation of primary human CD3^+^ T cells. In the presence of PD-L1, cTAC4.0 treatment at 10 nM and 100 nM significantly increased CD25 expression (**Fig. 3d-e**) and, to a lesser extent, CD69 expression (**Fig. S4c**) in T cells from two healthy donors. In contrast, CD80-Fc combined with anti-PD-L1-Fc or IgG4-Fc alone elicited minimal activation (**Fig. 3d-e** and **S4c**). Consistent with these results, cTAC4.0 enhanced T cell proliferation, as shown by an elevated percentage of cells undergoing two (D2) or three (D3) rounds of division, a reduced proportion of undivided cells (D0) (**Fig. 3f** and **S4d**), and an overall higher number of divided cells (**Fig. 3g**). Collectively, these findings demonstrated that cTAC4.0 effectively reversed PD-L1-mediated suppression, thereby promoting robust T cell activation and proliferation.

### cTAC4.0 primarily induced PD-L1 TE through CTLA-4, while triggering cell activation via CD28

In the presence of PD-L1-expressing target cells, cTAC4.0 simultaneously induced robust PD-L1 TE and T cell activation. To define the specific contributions of CTLA-4 and CD28 in these processes and test the performance of cTAC4.0 in different contexts, we engineered recipient Jurkat cells to express neither, one, or both receptors (**Fig. 4a**). cTAC4.0-mediated PD-L1 TE was most pronounced in CD28⁺CTLA-4⁺ Jurkat cells and, to a lesser extent, in CD28⁻CTLA-4⁺ Jurkat cells (**Fig. 4b**). Interestingly, a low but significant level of PD-L1 TE was also observed in CD28⁺CTLA-4⁻ Jurkat cells treated with cTAC4.0, whereas CD28⁻CTLA-4⁻ Jurkat cells showed no PD-L1 TE under any condition (**Fig. 4b**). These results indicated that CTLA-4 was the principal mediator of cTAC4.0-induced PD-L1 TE, with CD28 playing a minor contributory role.

**Figure 4.**
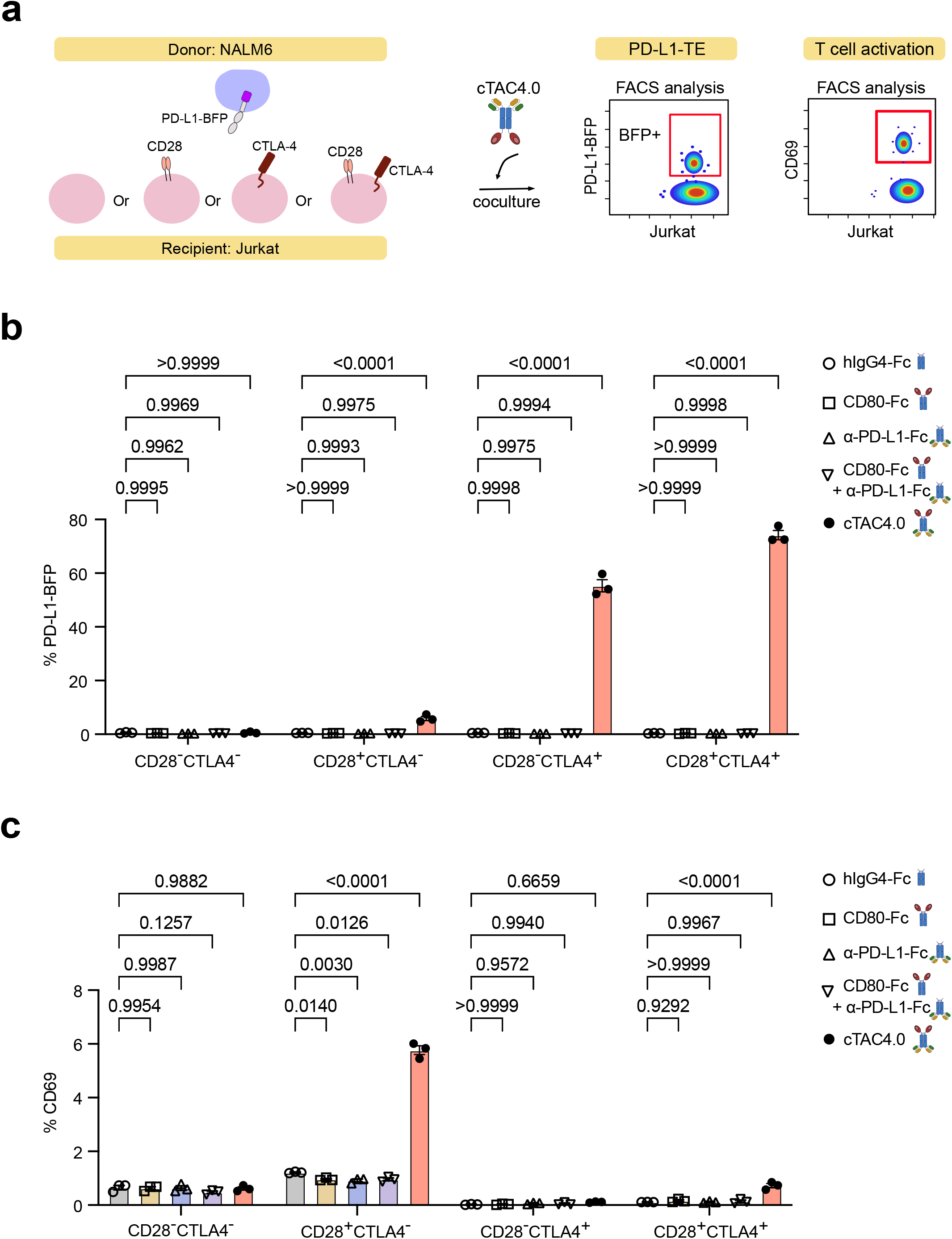
cTAC4.0 induces PD-L1 TE via CTLA-4 while promoting cell activation through CD28. **(a)** Schematics of workflow used in **b** and **c**. In **b**, cTAC mediated PD-L1 TE was quantified by BFP transfer from donor NALM6 cells expressing PD-L1-BFP to recipient Jurkat cells using flow cytometry. In **c**, cTAC4.0-mediated activation of Jurkat cells was measured by CD69 expression levels using flow cytometry. Four types of recipient Jurkat cells were utilized, each expressing either none, one, or both of CD28 and CTLA-4. **(b)** Quantification of transferred PD-L1-BFP on recipient Jurkat cells induced by different forms of antibodies at a concentration of 1nM. **(c)** Quantification of CD69 expression on recipient Jurkat cells by different forms of antibodies at a concentration of 1nM. Data are shown as mean ± s.e.m. Two-way ANOVA with Tukey’s multiple-comparisons test was used to assess significance (**b, c**). Exact P values are labeled above all comparisons. All numbers defined by ‘n’ indicate the number of biologically independent samples. Data are representative of two independent experiments.

We next assessed T cell activation by measuring CD69 expression on the same Jurkat cells (**Fig. 4a**). We found that cTAC4.0-mediated T cell activation was the highest in CD28⁺CTLA-4^-^ Jurkat cells and, to a lesser extent, in CD28^+^CTLA-4⁺ Jurkat cells, whereas CD28⁻CTLA-4⁺ and CD28⁻CTLA-4⁻ Jurkat cells exhibited minimal activation (**Fig. 4c**). These results demonstrated that cTAC4.0 required CD28 engagement to promote T cell activation. The reduced activation observed in CD28⁺CTLA-4⁺ Jurkat cells relative to CD28⁺CTLA-4⁻ Jurkat cells likely reflects the intrinsic inhibitory function of CTLA-4 ^24, 25^, preferential binding of cTAC4.0 to CTLA-4 over CD28 due to its higher affinity, or a combination of both. Collectively, these results demonstrated that CTLA-4 predominantly drove cTAC4.0-induced PD-L1 TE, whereas CD28 mediated cTAC4.0-induced T cell activation in the presence of PD-L1.

### cTAC4.0 mediated soluble PD-L1 degradation

Soluble PD-L1 (sPD-L1) is associated with poor prognosis in multiple solid tumors ^9, 10, 26^ and retains immune suppressive function comparable to its membrane-bound counterpart ^27^, making it a clinically relevant therapeutic target. To assess whether cTAC4.0 could facilitate sPD-L1 degradation, we incubated fluorescent dye labeled recombinant human PD-L1 (PD-L1-AF647) with CTLA-4^+^ Jurkat cells in the presence of cTAC4.0 or controls (**Fig. 5a**). cTAC4.0 induced a dose-dependent increase in PD-L1-AF647 uptake, whereas CD80-Fc, anti-PD-L1-Fc, their combination, or IgG4-Fc alone had minimal effect (**Fig. 5b-c**). To determine whether sPD-L1 was internalized or merely tethered to the cell surface, we examined its subcellular localization using AF488-labeled sPD-L1. The majority of AF488-labeled sPD-L1was detected in the cytosol with minimal colocalization with the membrane marker CD45 (**Fig. 5d**), indicating that cTAC4.0 primarily mediated sPD-L1 internalization. To evaluate lysosomal involvement in sPD-L1 degradation, we pretreated Jurkat cells with BafA1 and observed a small but significant increase in PD-L1-AF488 accumulation compared to DMSO-treated controls in the presence of cTAC4.0. (**Fig. 5e-f**). This increase was smaller than that observed for membrane-bound PD-L1 in **Fig. 2c-d**, likely reflecting the greater chemical stability of fluorophores such as AF488 relative to fluorescent proteins like BFP in the acidic lysosomal environment. Collectively, these results demonstrated that cTAC4.0 facilitated the internalization and lysosomal degradation of soluble PD-L1, highlighting its potential to target soluble immune suppressors and remodel the tumor microenvironment.

**Figure 5.**
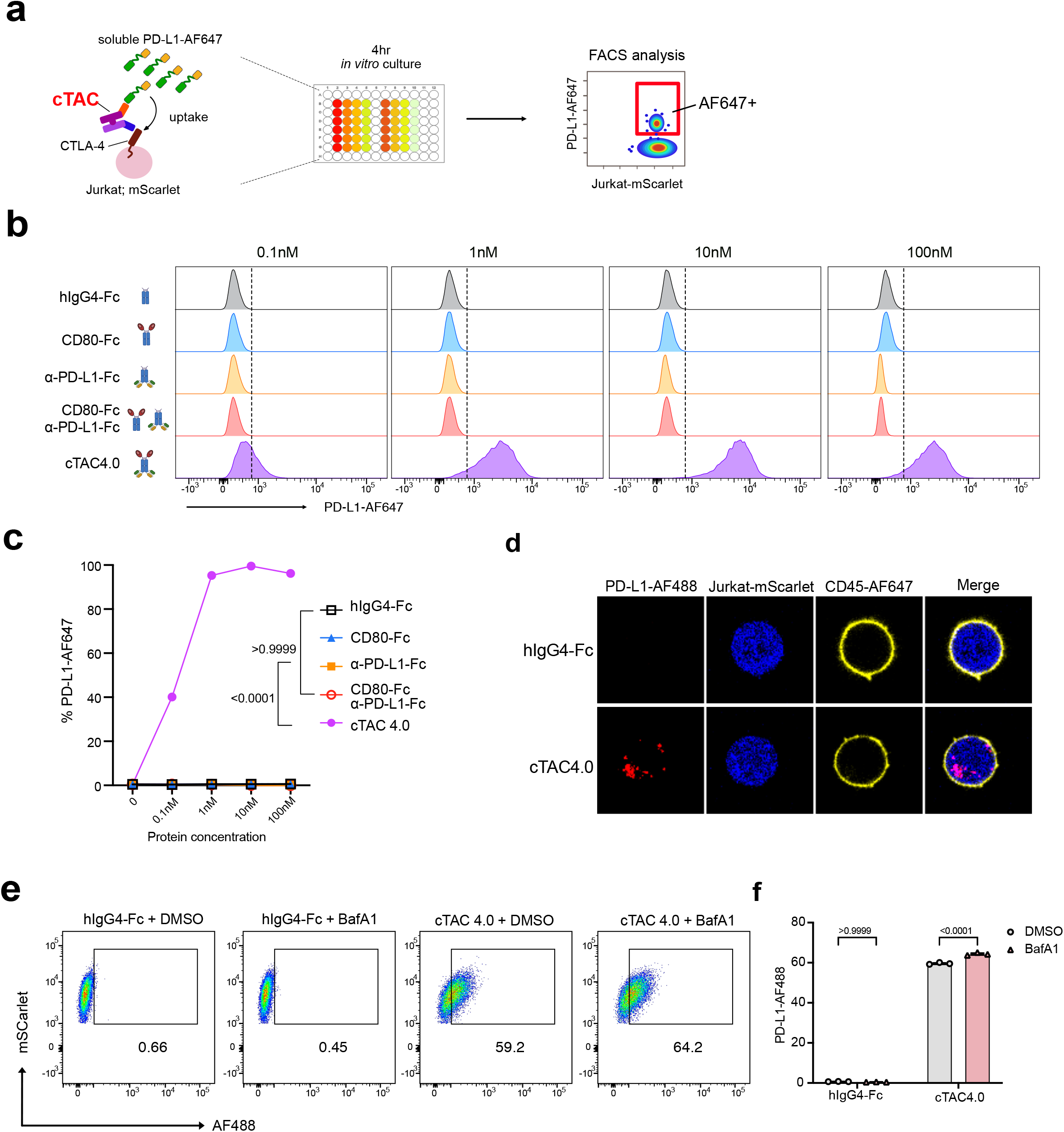
cTAC4.0 mediated free soluble PD-L1 uptake. **(a)** Schematics of workflow used in **b** and **c**. Soluble PD-L1 labeled with Alexa-Fluor647 (PD-L1-AF647) and cTAC at various concentrations were added in CTLA-4+ Jurkat cultures for 24hrs. Soluble PD-L1 uptake by Jurkat cells was measured by AF647 signals using flow cytometry. **(b)** Representative flow plots showing the intensity of PD-L1-AF647 uptake by recipient Jurkat cells at various concentration of antibodies. The gray dashed line represents the gating used to determine the percentages of AF647+ Jurkat cells, which are labeled as numbers on the plots. **(c)** Quantification of PD-L1-AF647 uptake by recipient Jurkat cells stimulated with cTAC4.0 at various concentrations. **(d)** Representative confocal images, demonstrating the subcellular distribution of free PD-L1 labeled with Alexa-Fluor 488 (PD-L1-AF488) in recipient Jurkat cells after incubating with 1nM of cTAC4.0 or controls for 24 hours. Cell membrane was indicated by surface staining of CD45 using Alexa-fluor 647 labeled antibody (CD45-AF647). **(e)** Representative flow cytometry plots to quantify the uptake of free PD-L1-AF488 in CTLA-4^+^ Jurkat cells induced by 0.1nM of cTAC4.0. Jurkat cells were pretreated with either DMSO or BafA1. **(f)** Quantification of the uptake of free PD-L1-AF488 in CTLA-4^+^ Jurkat cells in **e**. Data are shown as mean ± s.e.m. Two-way ANOVA with Tukey’s multiple-comparisons test was used to assess significance (**c, f**). Exact P values are labeled above all comparisons. All numbers defined by ‘n’ indicate the number of biologically independent samples. Data are representative of two independent experiments.

### cTAC4.0 synergized with T cell-based immunotherapies against solid tumor

Bispecific T cell engagers (BTEs) and T cells engineered with human leukocyte antigen (HLA)-independent TCR (HIT) represent promising strategies for targeting solid tumors by redirecting T cells to tumor antigens while bypassing HLA restriction and defective antigen presentation ^28, 29^. However, the immunosuppressive TME, particularly PD-L1-mediated inhibition ^30^ and insufficient co-stimulation ^28, 31^, remains a significant barrier to achieving sustained anti-tumor responses. Given that cTAC4.0 facilitates PD-L1 degradation and converts PD1-PDL1-mediated suppression into CD28-driven activation, we examined whether cTAC4.0 could boost the efficacy of these T cell-based therapies.

We first tested this hypothesis using an IL13Rα2-CD3 BTE targeting WM1366, a melanoma cell line that expresses both IL13Rα2 and PD-L1 ^32^. Engagement of T cells with WM1366 cells via this BTE robustly induced inhibitory receptors, including CTLA-4 and PD-1, on both CD4^+^ and CD8^+^ T cells (**Fig. S5a**). Such upregulation may contribute to tumor relapse, as co-treatment with cTAC4.0 markedly improved control of tumor regrowth compared with either the BTE alone or combination treatments using CD80-Fc and anti-PD-L1-Fc (**Fig. S5b–f**). These findings demonstrated that cTAC4.0 synergizes with BTEs to overcome PD-L1-mediated immune suppression and sustain T cell cytotoxicity.

We next evaluated whether cTAC4.0 could similarly augment the function of HIT-engineered T cells targeting EGFRvIII, a tumor-specific antigen in solid tumors. To generate these cells, we inserted an HDR template into exon 1 of the *TRAC* locus via CRISPR knock-in (**Fig. 6a**). Successful engineering was confirmed by co-expression of CD3 and an N-terminal Flag tag on the HIT *a*-chain (**Fig. 6b**). EGFRvIII-HIT T cells effectively lysed EGFRvIII-expressing U87 glioblastoma cells *in vitro* (**Fig. 6c**). Consistent with previous reports, repeated stimulation of EGFRvIII-HIT engineered T cells led to upregulation of multiple co-inhibitory molecules, including CTLA-4, PD-1, LAG-3, TIGIT, and TIM-3, particularly 24 hours after antigen exposure (**Fig. 6d**). Using EGFRvIII-HIT engineered T cells pre-stimulated with 0–2 rounds of U87 cells, we next examined whether cTAC4.0 could improve of cancer cell killing. Our data showed that cTAC4.0 significantly improved U87 killing by EGFRvIII-HIT T cells that had undergone one (1 pre-stim.) or two (2 pre-stim.) prior stimulations, particularly at low E/T ratios (**Fig. 6e**). In contrast, no enhancement was observed in fresh EGFRvIII-HIT T cells (0 pre-stim.), which expressed minimal CTLA-4 and PD-1 (**Fig. 6d-e**).

**Figure 6.**
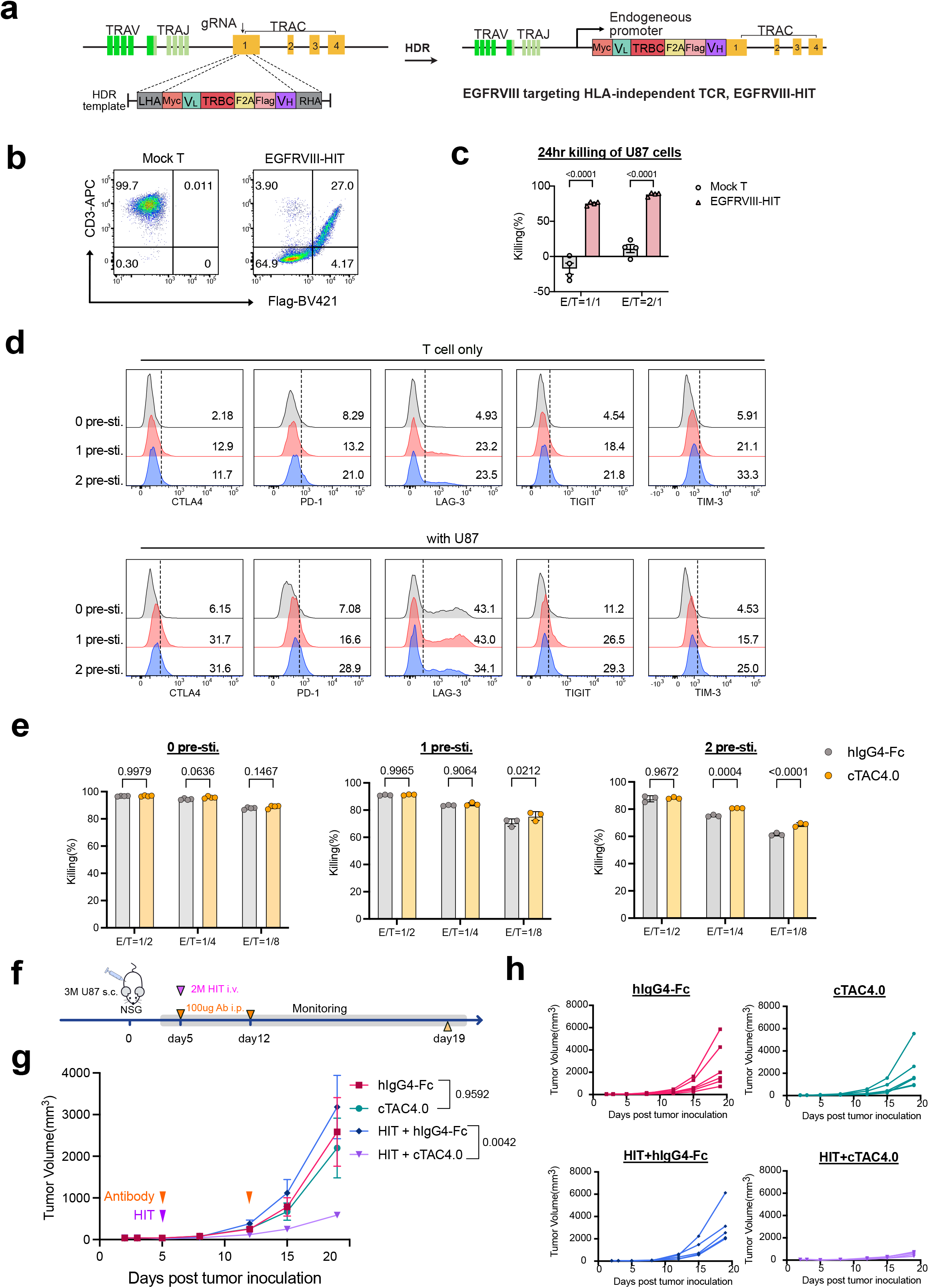
cTAC4.0 synergized with TCR-engineered therapeutic T cell against solid tumor. **(a)** Schematic diagram of CRISPR knock-in strategy to generate EGFRVIII targeting HLA-independent TCR, enables the expression of EGFRVIII-HIT using endogenous *TRAC* promoter. The variable fragments (V_L_ and V_H_) were developed from an EGFRvIII-specific antibody clone (mAb139). **(b)** Representative flow cytometry plots showing the generation of EGFRVIII-HIT, as indicated by the co-expression of CD3 and flag tag. **(c)** Quantification of 24-hour killing of U87 cells overexpressing EGFRVIII by EGFRVIII-HIT using a luciferase assay. **(d)** Representative flow plots showing the intensity of co-inhibitory molecules on EGFRVIII-HIT cells cocultured with (lower panel) or without (upper panel) U87 cells. EGFRVIII-HIT cells were previously stimulated with 0-2 rounds (corresponding to 0-2 pre-sti.) of U87 cells at an E to T ratio of 1:1, which is followed by one week resting. The gray dash line represents the gating used to determine the percentages of positive T cells, which are labeled as numbers on the plots. **(e)** Quantification of 24-hour killing of U87 cells overexpressing EGFRVIII by EGFRVIII-HIT cells, that were previously stimulated with 0-2 rounds (corresponding to 0-2 pre-sti.) of U87 cells as in **d**. (**f**) Schematic diagram of *in vivo* experimental workflow used in **g** and **h**, to test the synergistic effect between EGFRVIII-HIT and cTAC4.0 using U87 xenograft solid tumor model in NSG mice. **(g-h)** Tumor growth curves of U87 xenograft tumor, treated with/without EGFRVIII-HIT in combination with cTAC4.0 and the Fc control. Summarized and individual tumor growth data are presented in **g** and **h**, respectively. Data are shown as mean ± s.e.m. Two-way ANOVA with Tukey’s multiple-comparisons test was used to assess significance (**c, e, g**). Exact P values are labeled above all comparisons. All numbers defined by ‘n’ indicate the number of biologically independent samples. Data are representative of two independent experiments.

Finally, we evaluated the efficacy of cTAC4.0 *in vivo* using an EGFRvIII-U87 xenograft model (**Fig. 6f**). Neither cTAC4.0 alone nor EGFRvIII-HIT T cells with IgG4-Fc affected tumor growth relative to the control groups. However, combined treatment with cTAC4.0 and EGFRvIII-HIT T cells significantly suppressed tumor progression (**Fig. 6g-h**). These findings demonstrated that cTAC4.0 enhanced the anti-tumor activity of EGFRvIII-HIT engineered T cells by overcoming PD-L1-mediated suppression in the tumor microenvironment. Taken together, our findings established that cTAC4.0 effectively synergized with both BTEs and TCR-engineered T cells to overcome PD-L1-mediated immune suppression, thereby enhancing T cell function and anti-tumor efficacy both *in vitro* and *in vivo*.

## Discussion

The tumor microenvironment is highly immunosuppressive, driven by multiple mechanisms that fluctuate dynamically over time and across cell types^33, 34^, posing a critical challenge in restoring effective anti-tumor immunity. Recent immunotherapy strategies have focused on targeting individual immune checkpoints, such as PD-1/PD-L1, CTLA-4 or their combination^6^. However, these approaches benefit only a subset of patients, underscoring the need for innovative approaches that counteract immune suppression at multiple levels across diverse contexts. In this study, we demonstrated that cTAC4.0 achieves multi-level immune modulation through three synergistic mechanisms: (i) suppression of native CD80 TE via CTLA-4, thereby preserving costimulatory capacity on APCs; (ii) redirection of CTLA-4-mediated endocytosis to degrade PD-L1 *in trans*; and (iii) conversion of PD-L1 engagement into CD28-mediated co-stimulation. These unique features underscore cTAC4.0’s versatility in adapting to distinct immune environments. In suppressive tumor microenvironments where CTLA-4 expression predominates, cTAC4.0 can exploit CTLA-4-mediated TE to eliminate PD-L1 from opposing cells, thereby reducing inhibitory signaling. Conversely, in activated or co-stimulation-rich settings where CD28 engagement is more prominent, cTAC4.0 can bridge PD-L1⁺ target cells and CD28⁺ T cells to convert inhibitory cues into activating signals. Additionally, cTAC4.0 also demonstrates adaptability under diverse CD80 and PD-L1 expression profiles, mimicking the heterogeneity of the TME where PD-L1 and CD80 may be co-expressed on the same APCs or segregated between tumor and APCs. In both scenarios, cTAC4.0 efficiently removes PD-L1 while suppressing removal of CD80, which is required for costimulatory signaling. Together, this multi-functionality enables cTAC4.0 to maintain T cell activity across diverse physiological conditions, providing a dynamic means to sustain immune activation despite fluctuating receptor expression in the tumor microenvironment.

CTLA-4 naturally maintains immune homeostasis by capturing and internalizing costimulatory ligands from APCs ^14^, an evolutionarily conserved mechanism that limits T cell activation. We harnessed this intrinsic trafficking pathway to design a synthetic degradation system that functions *in trans*, directing CTLA-4-based internalization machinery toward selected targets such as PD-L1. To our knowledge, this represents the first demonstration that CTLA-4 TE can be redirected for targeted protein degradation, establishing a paradigm that merges immune checkpoint biology with degrader technology. Mechanistically, this strategy differs fundamentally from established approaches including PROTABs (utilizing surface E3 ligases) ^35^, TransTACs (utilizing transferrin receptor) ^12^, and LYTAC (utilizing mannose-6-phosphate receptor) ^36^, all of which rely on housekeeping processes within the same cell (*in cis*). In contrast, cTAC operates across cells, exploiting a physiological, immune-synapse-focused mechanism that becomes active primarily during T cell engagement. This design confers spatiotemporal precision and immune context specificity, potentially enabling more targeted immune regulation while minimizing systemic perturbations in healthy tissues.

The ability of cTAC4.0 to synergize with multiple T cell-based therapeutic modalities highlights its versatility as a platform technology. In the context of BTEs, cTAC4.0 alleviated adaptive suppression induced by prolonged T cell engagement, restoring cytotoxicity against PD-L1-expressing tumor cells. When combined with TCR-engineered T cells, cTAC4.0 enhanced effector function under conditions of repeated antigen exposure, which typically drive exhaustion. These findings indicate that cTACs function as modular immune amplifiers that can be layered onto diverse therapeutic platforms, from BTEs to TCR-engineered T cells. By coupling degradation to synapse formation rather than antigen identity, cTACs provide a unifying mechanism to overcome ligand-mediated immune resistance across tumor types, independent of the targeting modality employed.

Conceptually, cTACs represent a shift from signal blockade to synapse reprogramming. Conventional immunotherapies aim to block suppressive interactions, whereas cTACs reprogram them, transforming negative checkpoints into positive signals. This principle of functional inversion extends beyond PD-L1, providing a generalizable framework for engineering trans-cellular communications. Future iterations could target other suppressive ligand-receptor pairs, such as Galectin-9-TIM-3 ^37^ or CD47-SIRPα ^38^, potentially enabling a modular library of immune-restricted degraders that selectively modulate distinct immunosuppressive axes. Furthermore, integrating cTAC architectures with synthetic receptor circuits or next-generation receptor platforms could allow programmable control over degradation kinetics, co-stimulation, and effector functions. Synthetic receptor systems have demonstrated that T cells can execute logic-gated activation programs in response to specific antigen or environmental inputs ^39, 40^. Incorporating these principles into cTAC platforms could couple ligand degradation with T-cell signaling states, allowing degradation and activation to be dynamically coordinated within a single synthetic immune circuit. Such integration would position cTAC as a modular component that bridges molecular degradation with programmable cellular behavior.

Taken together, our findings establish cTAC as a first-in-class platform that co-opts natural immune trafficking for therapeutic protein degradation. By coupling checkpoint ligand elimination with T cell co-stimulation in an immune-synapse-enriched manner, this work establishes a paradigm for harnessing endogenous intercellular mechanisms to overcome resistance in cancer immunotherapy and provides a framework for engineering next-generation immune modulators that function at the tumor-immune interface.

## Methods

### Plasmid construction

The plasmids for the c-TAC bispecific antibodies were constructed using standard cloning methods as described previously ^41^. Briefly, DNA fragments encoding the human CD80 extracellular domain, a high affinity version of human PD-1 extracellular domain, anti-PD-L1-scFv (avelumab), and anti-CTLA-4-scFv (tremelimumab) were synthesized using human preferred codons and subcloned into an engineered pFuse-hIgG4 (S228P) vector for mammalian expression. The S228P mutation was introduced to prevent IgG4 Fab exchange ^42^. The plasmids used for producing stable cell lines were constructed with a second-generation lentiviral backbone previously described ^43^. Full length human PD-L1, CTLA-4, CD80 cDNA were ordered from Sino Bio, amplified, and cloned into the backbone. The NF-kB reporter plasmid backbone (pSIRV) was purchased from Addgene (Plasmid #118094) ^44^, in which the ECFP was replaced with a msfGFP, followed by an EFS promoter to drive the expression of puromycin resistance for selection.

### Cell culture

The HEK239T, NALM6, Jurkat, and U87 cells were purchased from the ATCC. Human peripheral blood mononuclear cells (PBMCs) were purchased from either STEMCELL Technologies or the Human Immunology Core of the University of Pennsylvania. U87 cells were engineered to express GFP, luciferase (GL) and EGFRvIII previously in lab ^45^. Human melanoma cell line WM1366 was acquired from the David B. Weiner lab (The Wistar Institute). The HEK239T, U87GL-EGFRvIII and WM1366 cells were cultured in DMEM (Gibco) with 10% FBS (Corning) and 200 U/ml penicillin–streptomycin (Gibco). NALM6 cells and Jurkat cells were cultured in RPMI-1640 (Gibco) medium supplemented with 10% FBS and 200 U/ml penicillin–streptomycin. Human PBMCs were cultured in X-VIVO 15 medium (Lonza) supplied with 5% human AB serum (MP Biomedical) and 10 ng/ml human IL-2 (Peprotech). All cells were grown at 37°C and 5% CO2 with saturating humidity, unless otherwise specified.

### Lentivirus, retrovirus and AAV production

A total of 40 million 293T cells (passage number <20) were seeded onto a 150-mm culture dish. The next day, the medium was replaced with pre-warmed fresh complete DMEM medium. For lentiviral production, 20 µg transfer plasmid, 10 µg psPAX2 (Addgene # 12260) and 5 µg pMD2.G (Addgene # 12259) were transfected with LipoD293 (SignaGen) following manufacturer’s instructions. For retroviral production, 16 μg pSIRV transgene vector plasmid and 8 μg pCL-Ampho (Novus Bio) were transfected with LipoD293. For both lentivirus and retrovirus production, supernatant was collected 48 hours post transfection, spun down at 3,000g for 15 minutes at 4°C, concentrated by adding 40% (w/v) PEG 8000 directly to the supernatant to final concentration of 8% (w/v) PEG 8000 and then incubated at 4 °C overnight. Viruses were then spun down at 1,500g for 30 minutes 4°C, and resuspended with complete DMEM or RPMI medium. For AAV production, 293T cells were transfected with AAV6 serotype plasmid, packaging plasmid pDF6, and transgene vector plasmid at a ratio of 1.7:2:1. 72 hours post-transfection, AAV particles were purified from the cell lysate using a previously described method ^46^. Briefly, 3 mL of pure chloroform was added to 30 mL of cell lysate, followed by vortexing for 5 minutes. Subsequently, 7.6 mL of 5 M NaCl was added, and the mixture was centrifuged at 3,000 × g for 5 minutes. The aqueous phase was collected and mixed with 9.4 mL of 50% (v/v) PEG 8000, followed by incubation on ice for 1 hour. The mixture was centrifuged at 3,000 × g for 30 minutes, and the supernatant was discarded. The viral pellet was resuspended in Dulbecco’s PBS supplemented with MgCl_2_. The viral solution was treated with universal nuclease (Thermo Fisher Scientific) at 37°C for 30 minutes, mixed with an equal volume of chloroform, vortexed, and centrifuged at 3,000 × g at 4°C for 5 minutes. The aqueous layer was collected and concentrated using a 100-kDa molecular weight cutoff filter (Millipore) by centrifugation at 3,000 × g. AAV viral titers were determined by quantitative PCR using custom TaqMan probes targeting the AAV inverted terminal repeats (ITRs). All viruses were aliquoted and stored at −80°C before usage.

### Stable cell line generation

NALM6 cells expressing PD-L1-BFP were generated using a lentiviral vector expressing human PD-L1 fused to mTagBFP2 via a GGGGS linker. NALM6-CD80-GFP cells were produced by replacing the PD-L1-GGGGS-mTagBFP2 construct with a CD80-GGGGS-eGFP construct. Finally, NALM6-PD-L1-BFP; CD80-GFP cells were generated by co-transducing NALM6 cells with the two aforementioned constructs. Parental Jurkat cells, which are CD28^+^CTLA-4^-^, were modified to generate Jurkat cell lines with distinct surface receptor profiles. CD28^+^CTLA-4^+^ Jurkat cells were created by transducing parental Jurkat cells with a lentiviral vector expressing human CTLA-4 followed by an mScarlet reporter, separated by a 2A sequence. To generate CD28^-^CTLA-4^-^ and CD28^-^CTLA-4^+^ Jurkat cells, CD28 was disrupted in parental Jurkat cells and CTLA-4-overexpressing Jurkat cells, respectively, using Cas9 ribonucleoprotein (RNP) electroporation with the Neon transfection system (Thermo Fisher). PD-1^wt^ CTLA-4^-^ and PD-1^trunc.^ CTLA-4^-^ Jurkat cells were produced by transducing parental Jurkat cells with lentiviral vectors expressing either full-length human PD-1 or a truncated version lacking the intracellular signaling domain, each paired with an mScarlet reporter separated by a 2A sequence. Similarly, PD-1^wt^ CTLA-4^+^ and PD-1^trunc.^ CTLA-4^+^ Jurkat cells were generated by transducing CTLA-4-expressing Jurkat cells with the same PD-1 constructs. To introduce an NF-κB reporter, Jurkat cells were transduced with a lentiviral vector expressing eGFP under the control of an NF-κB promoter, followed by an EFS promoter driving puromycin resistance for selection. All stable cell lines were purified based on reporter gene (GFP, BFP or mScarlet) expression using flow sorting or puromycin selection.

### Antibody production and purification

cTAC4.0, KIH-cTACs (1.0-3.0) ^47^, controls, and IL13Ra2-CD3 BTE were produced by transient transfection of Expi293F cells (Thermo Fisher, A14527) using the Expi293 Expression System Kit (Thermo Fisher, A14635). Expi293F cells were maintained in Expi293 Expression Medium at 37 °C with 8% CO_2_ on an orbital shaker at 120 rpm. When the cell density reached 4–6 × 10⁶ cells/mL, cultures were diluted to 3 × 10⁶ cells/mL prior to transfection. Plasmid DNA (1 µg per mL of culture) and ExpiFectamine 293 reagent were each diluted in Opti-MEM I Reduced Serum Medium, incubated for 5 min, and then combined and incubated for an additional 15 min to allow complex formation. The DNA–lipid complexes were slowly added to the prepared cells. After 18–22 h post-transfection, ExpiFectamine 293 Transfection Enhancer 1 and Enhancer 2 were mixed and gently added to the cultures according to the manufacturer’s instructions. Culture supernatants were harvested 6–7 days post-transfection, clarified by centrifugation, and filtered through a 0.22 µm membrane. The cTAC proteins were purified from the supernatants using Protein G affinity chromatography. Pierce disposable columns (Thermo Fisher, 29924) were packed with 2 mL of Pierce Protein G agarose resin (Thermo Fisher, 20399) and equilibrated with 5 mL of Protein G IgG Binding Buffer (Thermo Fisher, 21011). The diluted supernatant (1:1 with binding buffer) was incubated with the resin overnight at 4 °C with gentle stirring, then loaded onto the column. The column was washed with 15 mL of binding buffer, and bound proteins were eluted with 8 mL of IgG Elution Buffer (Thermo Fisher, 21004) in 1 mL fractions, which were immediately neutralized with 1 M Tris, pH 8.0 (Thermo Fisher, 15568025). Protein concentrations were determined by absorbance at 280 nm, and fractions containing the target protein were pooled. The buffer was exchanged into PBS using a Spectrum dialysis membrane (Fisher, 08-670-3BB). The IL13Ra2-CD3 BTE was purified using Ni-NTA affinity chromatography (Ni-NTA agarose resin, Qiagen) according to the manufacturer’s instructions as described previously ^32^. The final protein concentration was quantified using the Pierce BCA Protein Assay Kit (Thermo Fisher, 23225). The purity of all cTACs and controls were assessed by SDS-PAGE electrophoresis.

### Biolayer interferometry

All BLI experiments were conducted on an Octet RED96 (ForteBio) as described previously ^48^. To detect the binding affinity, serial dilutions of cTACs and controls were prepared as follows: CD28, 500nM to 15.625nM, 2-fold dilution; CTLA-4, 25nM to 0.78125nM, 2-fold dilution; PD-L1, 50nM to 1.5625nM, 2-fold dilution. Then, 25 ng/μL of His-tagged monomeric wild type CD28 extracellular domains (Sino Biological, 90182-C08H), or CTLA-4 extracellular domains (Sino Biological, 11159-H08H), or PD-L1 extracellular domains (Sino Biological, 10084-H08H) were immobilized onto optically transparent HIS1K biosensors (ForteBio). After washing with kinetic buffer, antigen-loaded sensors were subjected and titrated with each diluted concentration of homodimer cTAC and controls, respectively, as the analytes for hassociation and dissociation measurements. Dissociation constants (*K*_d_) and kinetic constants (*K*_on_ and *K*_off_) were calculated with the Octet Analysis software.

### Flow cytometry

All antibodies used in this study are listed in the Antibodies section of the Nature Portfolio Reporting Summary. Cell surface antigens were stained with MACS buffer (PBS with 0.5% BSA and 2mM EDTA) on ice for 15 minutes. LIVE/DEAD Fixable Near-IR Dead Cell Stain (Invitrogen) was included for primary T cell staining to exclude unspecific staining of dead cells. Samples were collected or sorted on a four-laser (405 nm, 488 nm, 561 nm and 640 nm) Aria II cell sorter (BD), five-laser (355 nm, 405 nm, 488 nm, 561 nm and 640 nm) Symphony S6 (BD), or FACSymphony A5 SE (BD). For sorting experiments, sorting purity was checked after every sort to make certain it was higher than 95%. All flow cytometry data were analyzed using FlowJo software v.10.10.0 (BD).

### Human T cell isolation

Human CD3^+^ T cells were isolated from PBMC samples (STEMCELL) using magnetic positive selection. Briefly, 15 million PBMCs were labeled with anti-human CD3-biotin (BioLegend, 1:100 dilution) in MACS buffer at 4°C for 10 minutes. After washing with MACS buffer, cells were resuspended in 120µL MACS buffer and incubated with 30 µL anti-biotin beads (Miltenyi) at 4°C for 15 minutes. CD3^+^ T cells were then isolated using MACS LS Columns and MACS Separators (Miltenyi).

### Trans-endocytosis assays

To assess CD80 and PD-L1 trans-endocytosis (TE) mediated by cTACs, NALM6 cells were co-cultured with Jurkat cells at a 1:1 E/T ratio for 4 or 24 hours, as specified in the figure legend. cTACs or control antibodies were added at concentrations ranging from 0 to 100nM. CD80 and PD-L1 TE were quantified by flow cytometry, measuring the percentage of CD80-GFP and PD-L1-BFP on Jurkat cells. For lysosome inhibition experiments, Jurkat cells were pre-treated with 100nM BafA1 at 37°C for 1 hour, followed by three washes prior to co-culture.

### T cell activation assay using Jurkat cell line

To evaluate cTAC-mediated T cell activation, Jurkat cells with or without an NFκB-GFP reporter were co-cultured with NALM6-PD-L1-BFP donor cells at a 1:1 effector-to-target (E/T) ratio for 24 hours. cTACs or controls were added at concentrations of 0 to 100nM. 0.5μg/mL anti-CD3 (Biolegend) were added to provide TCR stimulation. T cell activation was assessed by measuring the percentage of CD69-positive cells (in Jurkat cells without the NF-κB reporter) or NF-κB-GFP-positive cells (in Jurkat cells with the NF-κB reporter) using flow cytometry.

### Primary T cell activation and proliferation assay

To assess cTAC-mediated activation of primary CD3 T cells, donor NALM6-PD-L1-BFP cells were treated with 25μg/mL Mitomycin C (STEMCELL) at 37°C for 1 hour, washed three times, and co-cultured with freshly isolated CD3 T cells in the presence of 0.5 μg/mL anti-CD3 (Biolegend). For T cell proliferation assays, CD3 T cells were pre-stained with 10 μM eFluor450 proliferation dye (Thermo Fisher) prior to co-culture. cTACs or control antibodies were added at concentrations of 10 or 100 nM. T cell activation was quantified after 24 hours by measuring CD25 and CD69 expression using flow cytometry. T cell proliferation was assessed 3 days post co-culture by eFluor450 dye dilution using flow cytometry.

### Soluble antigen uptake

Jurkat cells overexpressing CTLA-4-2A-mScarlet were used to evaluate soluble PD-L1 uptake. Human recombinant PD-L1 labeled with AlexaFluor488 or AlexaFluor647 (R&D) was added at 1µg/mL as the soluble antigen source. cTACs or controls were included at concentrations ranging from 0.1 to 100nM. Cells were incubated at 37°C for 24 hours, washed twice, and analyzed by flow cytometry to determine the percentage of AlexaFluor488 or AlexaFluor647-positive Jurkat cells.

### Confocal imaging and quantification

Recipient Jurkat cells that had performed TE of CD80-GFP and/or PD-L1-BFP were sorted using flow cytometry. Cells were then stained with AlexaFluor647 labeled anti-CD45 (BioLegend) before imaging. About 10,000 purified Jurkat cells were seeded onto a µ-slide eight-well glass-bottom chamber (Ibidi) that had been previously treated with poly-L-lysine (Sigma). Cells were imaged by Leica Stellaris 8 FALCON with the 40× objective (oil). Representative images were exported using ImageJ (https://imagej.nih.gov/). The percentage of GFP and BFP membrane colocalization was analyzed by a customized MATLAB (https://github.com/xs254/Colocalization_calculator/tree/main) (Mathworks, Natick, MA) described before ^43^. Antigen-membrane colocalization percentage was calculated by the following formula where *I_antigen-cytosol_*, *I_antigen-cell_* stands for intensity of each pixel classified as foreground in both antigen and cytosol mask, respectively. *I_antigen background_* stands for mean intensity of background pixels in antigen image (1.25 used for all analysis).

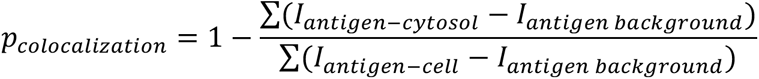

### Flow cytometry analysis of T-cell exhaustion following BTE treatment

To assess T-cell exhaustion phenotypes induced by IL13Rα2-CD3 BTE treatment, melanoma WM1366 cells were cocultured with peripheral blood mononuclear cells (PBMCs) from the indicated donors. WM1366 cells were seeded in 24-well plates at a density of 5 × 10⁴ cells per well and cultured for 24 hours prior to treatment. IL13Rα2-CD3 BTE was added to the wells at a final concentration of 300 ng/mL, followed by the addition of PBMCs at an effector-to-target (E:T) ratio of 10:1. After 96 hours of coculture, suspension cells were collected for flow cytometric analysis. For staining, cells were first incubated with APC anti-human CTLA-4 antibody (BioLegend, 349907) at 37 °C for 30 minutes, washed, and then stained at 4 °C for 30 minutes with the following antibody cocktail: Fixable Viability Dye eFluor 780 (Thermo, 65-0865-14), PerCP-Cy5.5 anti-human CD3 (BioLegend, 344808), FITC anti-human CD4 (BioLegend, 344604), Pacific Blue anti-human CD8 (BioLegend, 344718), and BV711 anti-human PD-1 (BD Biosciences, 564017).

### *In vitro* cytotoxicity analysis using xCELLigence RTCA eSight

To evaluate the in vitro efficacy of cTAC in combination with the IL13Rα2-CD3 BTE, a cell impedance–based real-time cytotoxicity assay was performed using the xCELLigence RTCA eSight system (Agilent Technologies). Melanoma WM1366 cells were seeded in an E-plate (Agilent, 300601010) at a density of 5,000 cells per well and cultured for 24 hours prior to treatment. IL13Rα2-CD3 BTE and cTAC proteins were diluted in complete medium and added to the wells at final concentrations of 300 ng/mL and 100 nM, respectively. Peripheral blood mononuclear cells (PBMCs) from the indicated donors were then added at an effector-to-target (E:T) ratio of 10:1. The real-time cell index and bright-field images were continuously recorded and analyzed using the RTCA Software (Agilent Technologies).

### TCR-engineered T cells

Purified CD3^+^ T cells were stimulated with Dynabeads Human T-Activator CD3/CD28 (Thermo Fisher Scientific) at a 1:1 bead-to-cell ratio for 48 hours. EGFRvIII-targeting TCR-engineered T cells were generated by disrupting the TRAC locus using the Cas9 ribonucleoprotein (RNP) method, followed by AAV6-mediated delivery of an HDR template. Briefly, chemically synthesized Cas9 crRNA and tracrRNA (IDT) were resuspended in nuclease-free IDTE buffer at 200 µM, mixed at a 1:1 ratio, and annealed as an sgRNA by heating to 95°C for 5 minutes and cooling to 37°C for 10 minutes. RNPs were formed by combining SpCas9 nuclease (IDT) with the annealed sgRNA at a 1:2 Cas9-to-sgRNA molar ratio and incubating at room temperature for 15 minutes. The RNPs were mixed with activated CD3+ T cells and electroporated using the Neon transfection system (1600 V, 10 ms, 3 pulses). Cells were transferred to prewarmed complete X-vivo medium (Lonza). Approximately 10¹¹ AAV6 viral particles were added to the culture post-electroporation. The knock-in efficiency was verified 6 days later by staining CD3 and the flag tag. CD3^+^ Flag^+^ EGFRvIII-targeting TCR-engineered T cells were sorted using Aria II flow sorter (BD).

### Luciferase-based cytotoxicity assay

The cytotoxicity of EGFRvIII-targeting TCR-engineered T cells was assessed using a luciferase-based assay. Luciferase- and EGFRvIII-expressing U87 cells served as targets. Effoctor T cells were pre-stimulated with 0-2 rounds of EGFRvIII⁺ U87 cells at an E: T ratio of 1:1, followed by a 3-day rest before assay. For cytotoxicity measurement, effector and target cells were co-cultured in triplicate at the indicated E:T ratios in white-walled, clear-bottom 96-well plates (Corning) with cTAC4.0 or the control antibody. After 24 hour, 150 μg/mL D-luciferin (PerkinElmer) was added directly to each well, and luminescence was measured after 5 min using a plate reader (PerkinElmer). Tumor cell killing was quantified by normalized luminescence relative to wells containing target cells alone.

### *In vivo* tumor modeling

NSG mice (Jackson Laboratory) were bred in-house and used for experiments. Female NSG mice (6–10 weeks old) were subcutaneously inoculated with 3 million U87GL-EGFRvIII cells on day 0. On day 5, 2 million EGFRvIII-targeting TCR-engineered T cells were injected intravenously. cTAC4.0 or controls (100 µg) were administered intraperitoneally on days 5 and 12. Tumor burden was monitored by measuring tumor volume, calculated using the formula: volume = (π/6) × length × width × depth.

## Acknowledgments

We thank all members of the Chen laboratory and the Zhou laboratory, as well as our colleagues at Yale and The Wistar Institute for assistance and/or discussion. We thank the various core facilities at Yale for technical support. We thank The Wistar Institute’s Flow Cytometry Facility and Research Supply Center for providing technical support. We thank Max Eldabbas, Emileigh Maddox, Tanishk Sinha, and Jiayi Shu of the Human Immunology Core at the Perelman School of Medicine at the University of Pennsylvania for providing de-identified, human PBMCs. The HIC is supported in part by NIH P30 AI045008 and P30 CA016520. S.C. is supported by NIH (R33CA281702), DoD (W81XWH-21-1-0514, HT94252310472), Cancer Research Institute (Lloyd J. Old STAR Award (CRI4964)), Sontag Foundation (DSA), Pershing Square Sohn Cancer Research Alliance, and Chen lab discretionary funds from Yale School of Medicine. X.Z. is supported by NIH/NCI Pathway to Independence Awards (K99/R00), with the K99 work performed at Yale University (K99CA282989) and the R00 work at The Wistar Institute (R00CA282989). K.T. is supported by Gruber Science Fellowship.

## Institutional Approval

This study has received institutional regulatory approval. All recombinant DNA and biosafety work were performed under the guidelines of Yale Environment, Health and Safety (EHS) Committee with approved protocols (Chen-rDNA 15-45), and The Wistar Institute’s Institutional Biosafety Committee (IBC) with approved protocol # 22504650. All human sample work was performed under the guidelines of Yale University Institutional Review Board (IRB) with an approved protocol (HIC#2000020784). All animal work was performed under the guidelines of Yale University Institutional Animal Care and Use Committee (IACUC) with approved protocols (Chen-20068).

## Statistics and Schematic illustrations

Statistics for standard biological assays were done using Prism, with detailed method indicated in Figure legends. Schematic illustrations were created with Affinity Designer.

## Data and materials availability

All data generated or analyzed during this study are provided. Source data and statistics are provided in an excel file.

---

**Figure S1.**
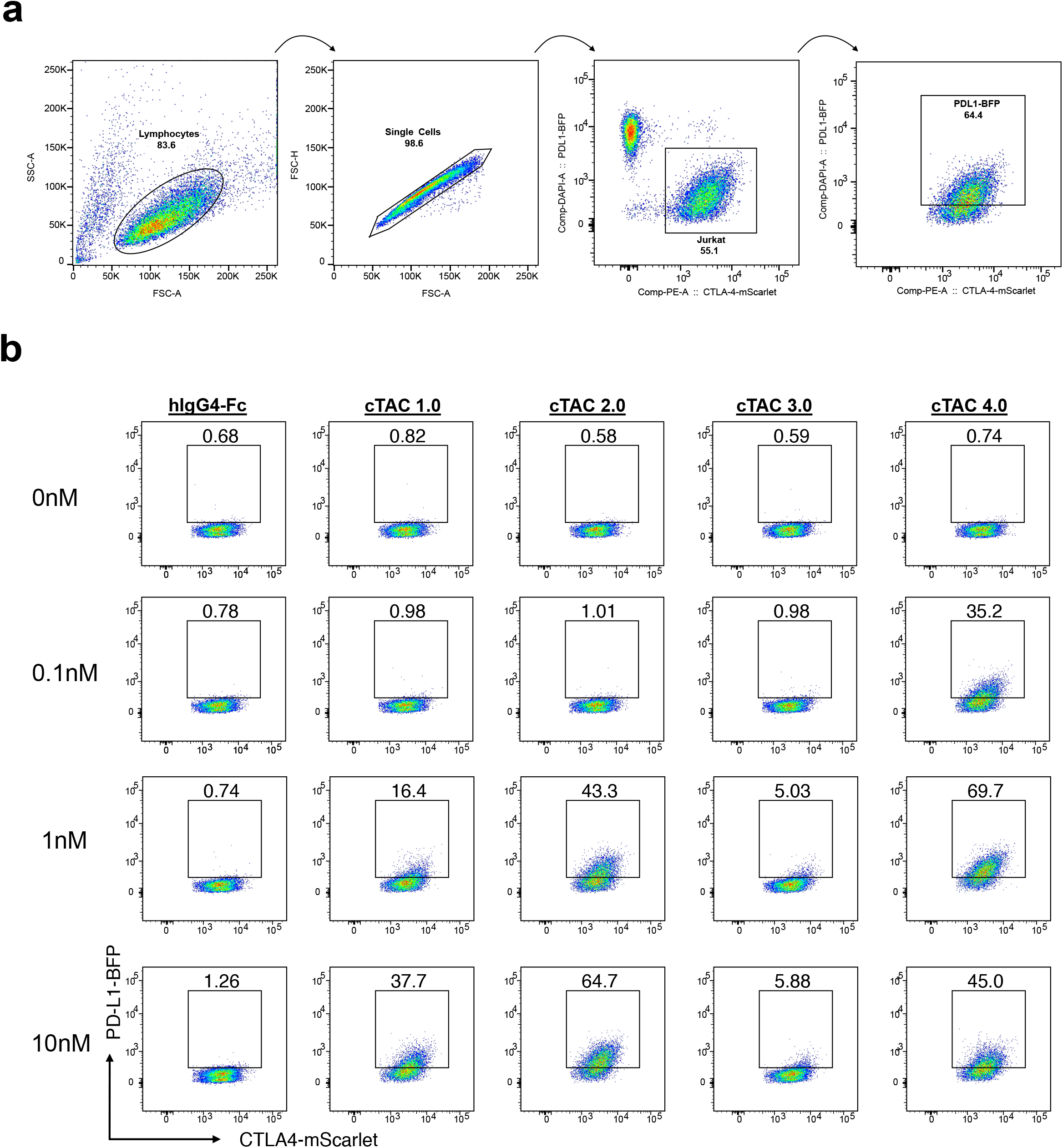
Comparisons of PD-L1 TE mediated by different cTACs. **(a)** Gating strategy used in b and Fig.1d. **(b)** Representative flow cytometry plots showing the percentage of PD-L1-BFP on recipient Jurkat cells after coculture with NALM6-PD-L1-BFP in the presence cTACs at various concentrations.

**Figure S2.**
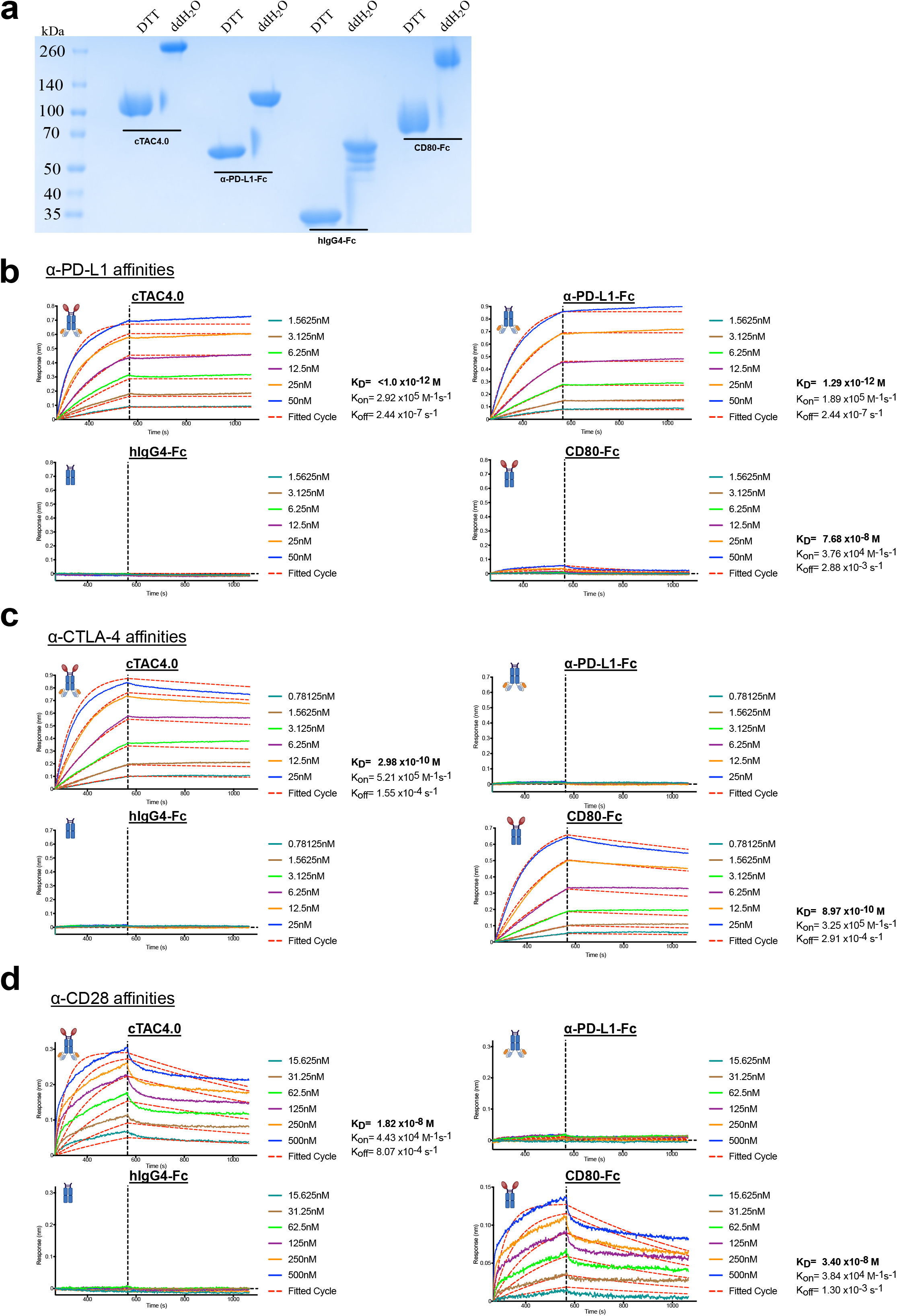
cTAC4.0 showed strong binding to human PD-L1, human CTLA-4, and relative weak binding to human CD28. **(a)** Coomassie-stained SDS-PAGE analysis of four purified resultant proteins. The protein samples were analyzed under reducing condition (+DTT) and nonreducing conditions (+ddH2O). **(b)** Binding affinities of cTAC4.0 and controls to monomeric wild type PD-L1 as measured by BLI. PD-L1(His Tag) recombinant protein was immobilized on an anti-Penta-His (HIS1K) sensor and tested for real-time association and dissociation of cTAC4.0 and controls. Global fit curves are shown as red dotted lines. The vertical dashed lines indicate the transition between the association and dissociation phase. **(c)** Binding affinities of cTAC4.0 and controls to monomeric wild type CTLA-4 as measured by BLI. CTLA-4 (His Tag) recombinant protein was immobilized on an anti-Penta-His (HIS1K) sensor and tested for real-time association and dissociation of cTAC4.0 and controls. Global fit curves are shown as red dotted lines. The vertical dashed lines indicate the transition between the association and dissociation phase. **(d)** Binding affinities of cTAC4.0 and controls to monomeric wild type CD28 as measured by BLI. CD28 (His Tag) recombinant protein was immobilized on an anti-Penta-His (HIS1K) sensor and tested for real-time association and dissociation of cTAC4.0 and controls. Global fit curves are shown as red dotted lines. The vertical dashed lines indicate the transition between the association and dissociation phase.

**Figure S3.**
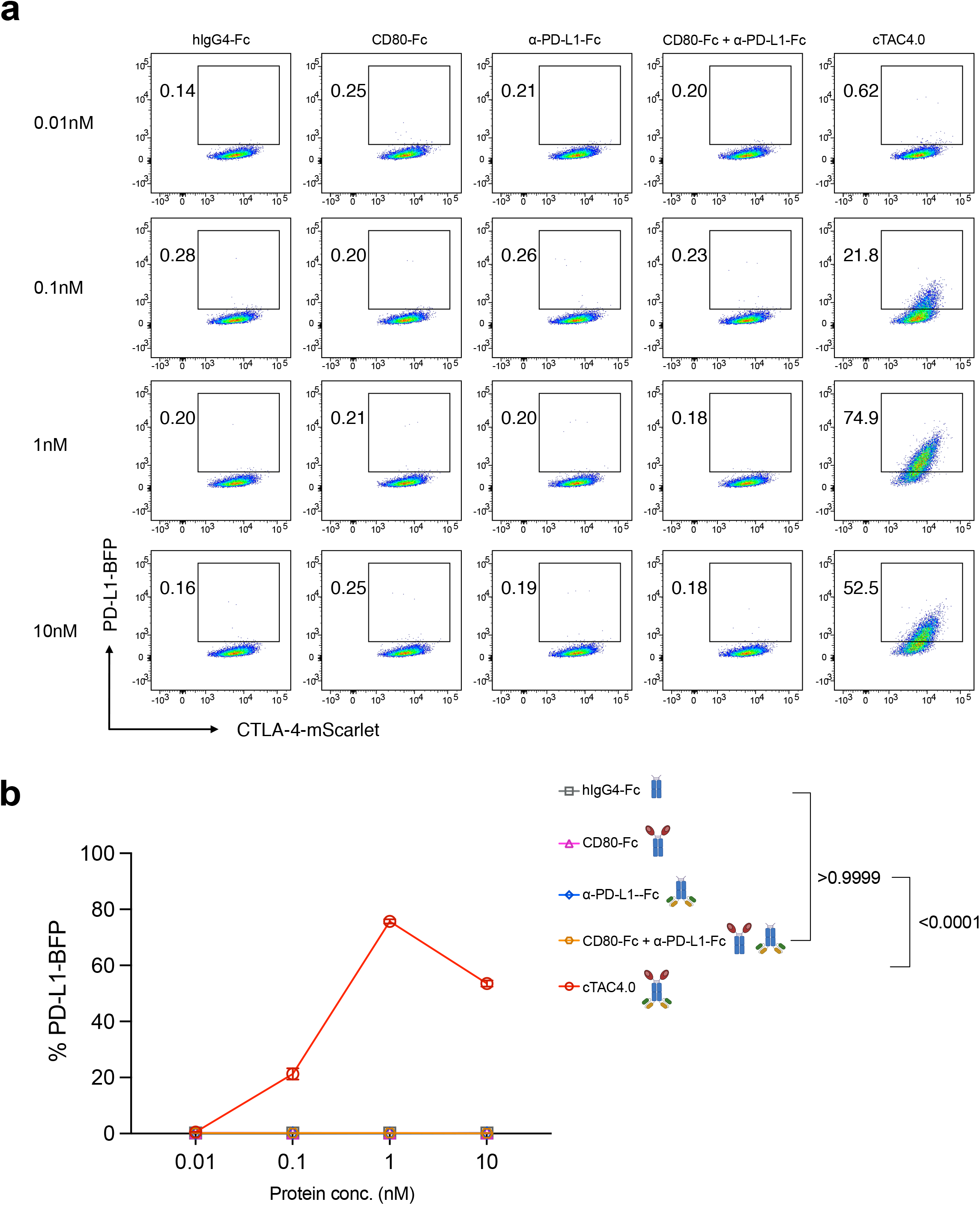
PD-L1 TE is induced by cTAC4.0, but not CD80-Fc, anti-PD-L1-Fc or the combination of those two antibodies. **(a)** Representative flow cytometry plots showing the percentage of PD-L1-BFP on recipient Jurkat cells after coculture with NALM6-PD-L1-BFP in the presence of cTAC4.0 and its control antibodies at various concentrations. **(b)** Quantification of **a**. Data are shown as mean ± s.e.m. Two-way ANOVA with Tukey’s multiple-comparisons test was used to assess significance (**b**). Exact P values are labeled above all comparisons. All numbers defined by ‘n’ indicate the number of biologically independent samples. Data are representative of two independent experiments.

**Figure S4.**
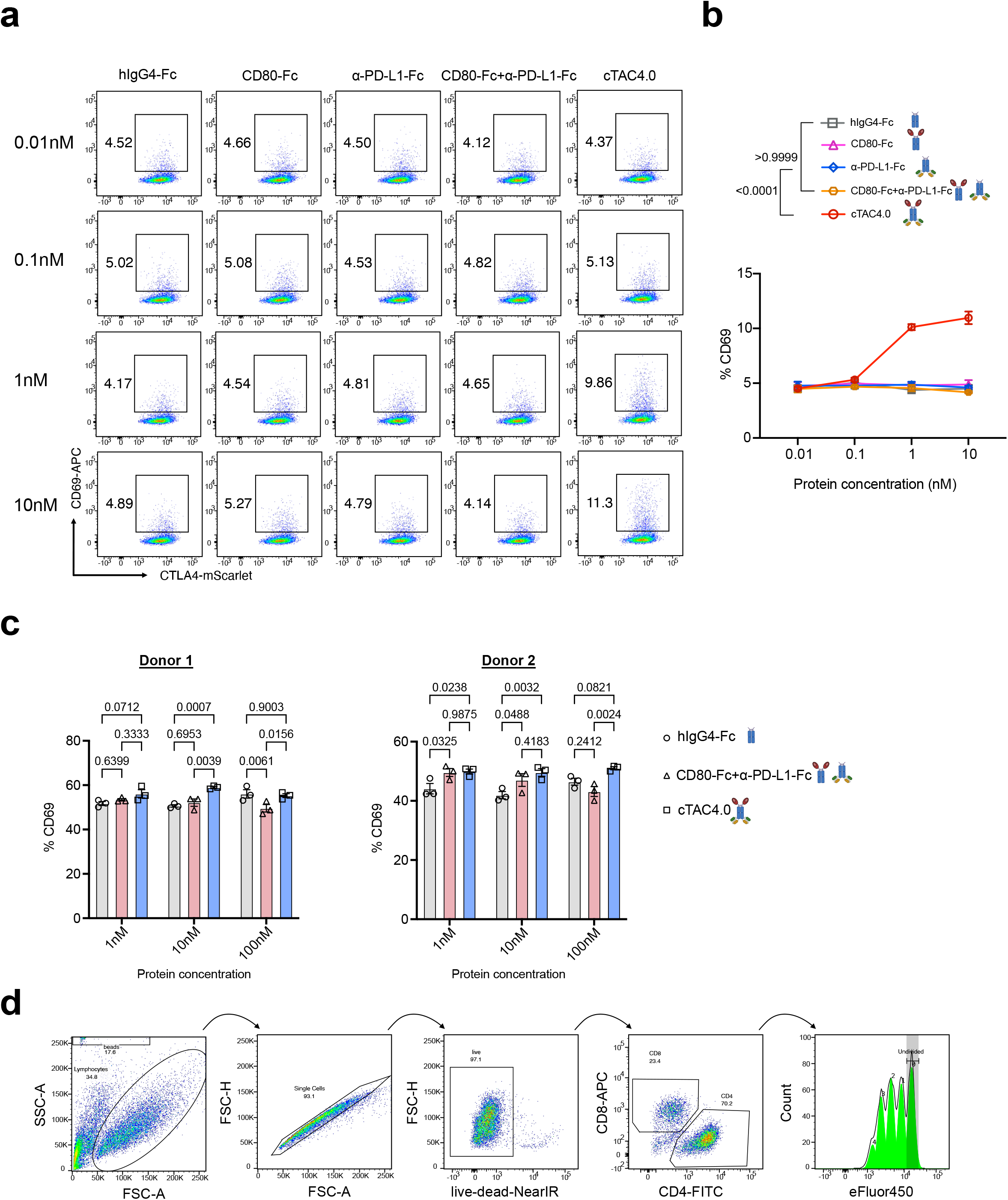
cTAC4.0 converted PD-L1-mediated suppression to T cell activation. **(a)** Representative flow cytometry plots showing the percentage of CD69 on recipient Jurkat cells after coculture with NALM6-PD-L1-BFP in the presence of cTAC4.0 and its control antibodies at various concentrations. **(b)** Quantification of **a**. **(c)** Quantification of T cell activation induced by varying concentrations of cTAC4.0 in CD3 T cells from two healthy donors, assessed by CD69 expression levels. **(d)** Representative flow cytometry plots showing the gating strategy used in **Fig. 3f-g**. Data are shown as mean ± s.e.m. Two-way ANOVA with Tukey’s multiple-comparisons test was used to assess significance (**b, c**). Exact P values are labeled above all comparisons. All numbers defined by ‘n indicate the number of biologically independent samples. Data are representative of two independent experiments.

**Figure S5.**
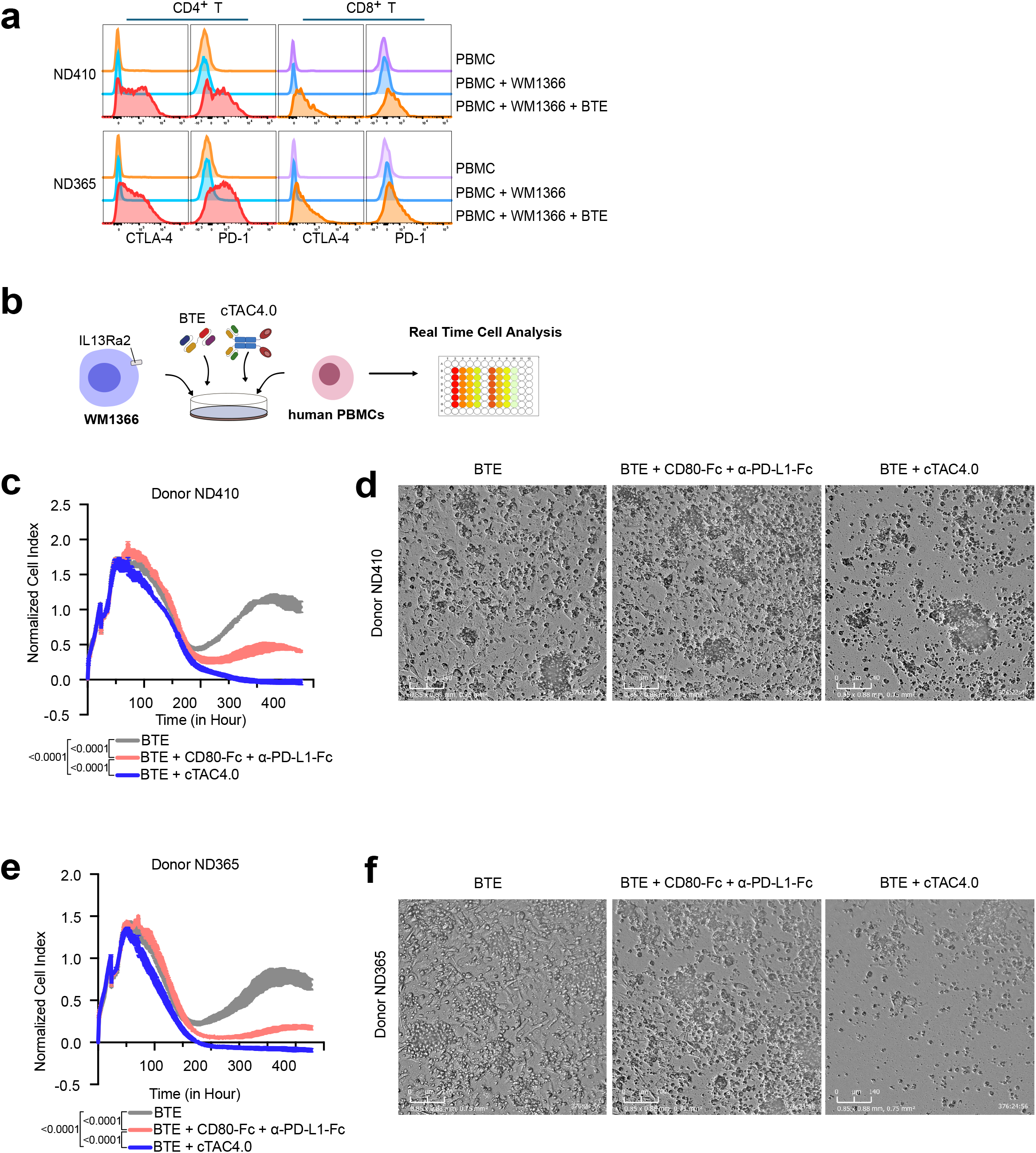
cTAC4.0 enhances the cytotoxicity of BTE *in vitro*. (**a**) Flow cytometry plots showing the upregulation of CTLA-4 and PD-1 on the CD4 and CD8 T cells within the PBMCs after coculturing with IL13Ra2-expressing WM1366 cell line and IL13Ra2-CD3 BTE. (**b**) Schematics of workflow used in **c-f**. IL13Ra2-expressing WM1366 cell line was cocultured with healthy human PBMCs in the presence of IL13Ra2-CD3 BTE and cTAC4.0. Real-time cell analysis was used to track the tumor lysis. (**c-f**) Cytotoxicity measured by xCELLigence RTCA assay to show the synergy of BTE and cTAC4.0 using PBMCs from two donors, as demonstrated by the normalized cell index (**c** and **e**), and bright-field images (**d** and **f**). Data in **c** and **e** are two technical replicates shown as mean ± s.e.m. Time-course data were first analyzed by two-way ANOVA to assess treatment effects across the entire curve. To compare overall responses between groups, area-under-the-curve (AUC) values were calculated and analyzed by one-way ANOVA, followed by Tukey’s test to correct for multiple comparisons. Data are representative of two independent experiments.

